# Isolation Methods of Tumor Endothelial Cells Impact AngiomiR Profiles

**DOI:** 10.1101/2025.06.25.661572

**Authors:** Lincy Edatt, Gargi Dixit, Salma H. Azam, Yihsuan S. Tsai, Andrew C. Dudley, Chad V. Pecot

**Affiliations:** Lineberger Comprehensive Cancer Center, University of North Carolina at Chapel Hill, Chapel Hill, NC 27599, USA; Department of Medicine, University of North Carolina at Chapel Hill, Chapel Hill, NC 27599, USA; Department of Microbiology, Immunology, and Cancer Biology, The University of Virginia, Charlottesville, VA 22908, USA; UVA Comprehensive Cancer Center, The University of Virginia, USA; RNA Discovery Center, University of North Carolina at Chapel Hill, Chapel Hill, NC 27599, USA

## Abstract

MicroRNAs (miRNAs) play an important role in endothelial cell growth and differentiation. Tumor angiogenesis-specific miRNAs (angiomiRs) are a subset of miRNAs that are dyregulated in tumor endothelial cells. Because of the importance of angiogenesis in cancer progression, regulation of angiomiRs may have significant therapeutic implications. However, discovery of angiomiRs has often been limited by biased model systems that may not be valid. Here, we evaluated whether the variable expression levels of angiomiRs in endothelial cells were impacted by the isolation methods used to profile them. Using an autochthonous, genetically engineered mouse model of lung adenocarcinoma, we used Nanostring to profile miRNA expression levels of normal lung endothelial cells (NECs) to tumor endothelial cells (TECs) using two endothelial cell (EC) isolation methods: 1) staining and sorting ECs directly from tumors (“*in vivo*”), and 2) magnetic bead isolation and sub-cloning ECs (“*in vitro*”). We then compared candidate angiomiRs with the profiles from two orthotopic, immunocompetent lung cancer models. When TECs were directly enriched from tumors (“*in vivo*” method), three candidate angiomiRs (miR-30b, miR-1981, and miR-707) were significantly lower in TECs than NECs. In contrast, when ECs were isolated and cultured (“*in vitro*” method), three different candidate angiomiRs (miR-200a, miR-124 and miR-186) were significantly lower in TECs than NECs. Using two independent model systems for validation, we found miR-30b to be significantly reduced in TECs using freshly sorted ECs. Conversely, the *in vitro* discovered angiomiR candidates did not validate in these model systems, suggesting that TECs grown *in vitro* may not maintain relevant angiomiR profiles or serve as an adequate method for molecular profiling. Our findings demonstrate that angiomiR expression patterns are impacted by isolation methods. Instead of relying on ECs cultured *in vitro*, we suggest careful validation studies of cells freshly collected from tumors before determining whether a miRNA is a bona fide angiomiR.

## INTRODUCTION

Micro-RNAs (miRNAs) are short noncoding RNAs that play important roles in cell growth and differentiation. These miRNAs impact cells during many metabolic processes including cell division^1^, cell stability^2^ and cell death^3^ through post-transcriptional regulation of mRNA by splicing and degradation^4^. It has been demonstrated that the expression of miRNAs also significantly contributes to the modulation of tumor microenvironments (TME)^5^ and can be both pro-angiogenic and anti-angiogenic^6,7^. Several miRNAs specific to tumor endothelial cells (TECs), termed angiomiRs, have been found to significantly impact the progression of vascularization and angiogenesis during tumor progression^8,9^. Pro-angiogenic miRNAs like miR-9, miR-138 and miR-494 induce angiogenesis in glioma demonstrating their conserved role in different cancer types^6^. The functions and regulatory mechanisms of miRNAs have been elucidated by employing various *in vitro, in vivo* and *in silico*, approaches. Unfortunately, if the chosen method of isolation of TECs does not recapitulate the tumor microenvironment, analyses of these identified miRNAs may not be relevant to the pathophysiology in the TME.

AngiomiRs have been widely studied employing two different approaches, *in vitro* cultured ECs or freshly isolated ECs, usually obtained using fluorescence activated cell sorting (FACS). Because sorting of ECs from freshly excised tumors is not always feasible, most published studies have relied on cultured systems to mimic EC conditions in the TME. Some studies have found similar miRNA expression profiles from both cultured and freshly sorted ECs, which has led to the increased use of cultured tumor-derived ECs to study angiomiRs^10^. However, others have observed that miRNA profiles from cultured cells can significantly vary from those obtained from freshly isolated cells^11^.

To study the discrepancies attributed to the expression levels of miRNAs by the differences in isolation methods, we performed a Nanostring-based profiling study. Using an autochthonous, genetically engineered lung cancer model, we compared expression levels of miRNAs between normal ECs (NECs) with TECs when directly sorted from lung cancer tumors, or when the ECs were enriched and isolated using a controlled cell culture environment. We then used several independent, syngeneic lung cancer model systems to validate candidate angiomiRs. Our findings demonstrate highly discordant miRNA levels in ECs based on isolation method, and suggests caution is needed when selecting candidate angiomiRs for further study.

## METHODS

### Isolation and purification of ECs

Endothelial cells were isolated from lung tumors after 8 weeks of tumor induction following intra-tracheal instillation of AdenoCre in a Lox-Stop-Lox (LSL)-KRAS^G12D^; Lkb1^L/L^; p53^L/L^ (KLP) mouse model^12^, as well as from the lungs of healthy, age-matched non-tumor bearing KLP control mice. The KLP mice were kindly provided by Dr. William Y. Kim at UNC. FACS was employed to isolate ECs using CD45, EPCAM, CD31 and LYVE1 markers as previously described^13^. *Cdh5-Cre*^*ERT2*^ were crossed with Ai6 ZSGreen reporter mice to facilitate EC isolation^14^. *Cdh5-Cre*^*ERT2*^ mice were generated by Ralf Adams (Max Planck Institute for Molecular Biomedicine).

In the ZsG mouse model systems LLC-fLuc and LN2A-fLuc, necropsies were performed after 14-20 days, and the tumors were harvested and dissociated at 4° in buffered collagenase^15^, stained with fixable dead cell stain and then sorted for live cells that are ZsG+

### Culturing ECs

For culturing normal lung and tumor-derived ECs, CD31+ cells isolated from lung tissue were purified using a magnetic separation technique and a-CD31 polyethylene (PE) conjugated to a-PE magnetic beads as described previously^15^. After purification, ECs were cultured in 0.01% D-glucose DMEM with 10% FBS, 10% NuSerum IV, 0.001%. vascular endothelial growth factor, 0.0005% *b*-fibroblast growth factor, and 0.1% porcine heparin. Clonal rings were used to isolate small culture colonies, and the selected clones were further propagated by passaging until cell health declined.

### *In Silico* Analyses of miRNAs using Nanostring

Expression levels of overall miRNAs from both normal endothelial cells (NECs) and tumor endothelial cells (TECs) from KLP mice were obtained using ROSALIND software after Nanostring’s nCounter Analysis System was performed on both freshly isolated and cultured Ecs to show the distribution of distinct miRNA populations by using a threshold log-fold change value of 1.5 and a p-value of 0.05. Volcano plots were used to show the fold change of individual miRNAs of TECs normalized to NECs on both freshly isolated and cultured ECs.

### RNA Isolation and qPCR for miRNAs

To quantitate miRNAs using qPCR from both freshly isolated ECs and cultured ECs, total miRNA was extracted from freshly isolated cells using the Invitrogen PureLink miRNA Isolation Kit based on the manufacturer’s protocol. cDNA was synthesized for the RNA fraction using Applied System’s TaqMan advanced miRNA cDNA synthesis kit as instructed by the manufacture. To selectively amplify candidate miRNAs, PCR and requisite primers were employed to amplify miR-30B, miR-200A, miR-124 and miR-186.

### Statistics

Data was analyzed using 2-tailed Student’s *t* test (for comparisons of 2 groups). *P* value less than 0.05 was deemed statistically significant. All statistical tests for in vitro and in vivo experiments were performed using GraphPad Prism 7 (GraphPad Software).

## RESULTS

### *Comparing In Vivo* and *In Vitro* Detection Methods

Using an autochthonous Lox-Stop-Lox (LSL)-KRAS^G12D^; Lkb1^L/L^; p53^L/L^ (KLP) mouse model, mice were intra-tracheally instilled with 10^6^ plaque forming units of adenoviral Cre recombinase and monitored by *in vivo* CT imaging for tumor growth at both 4 weeks and 8 weeks (Figure 1A), and age-matched control lungs (Figure 1B) and Adeno-Cre treated mice were imaged at 8 weeks for tumor excision (Figure 1C). Tumors from these mice were dissociated with collagenase and ECs were isolated using two different methods. First, using magnetic separation and a-CD31, cultured and clonally selected (Figure 1D).

**Figure 1:**
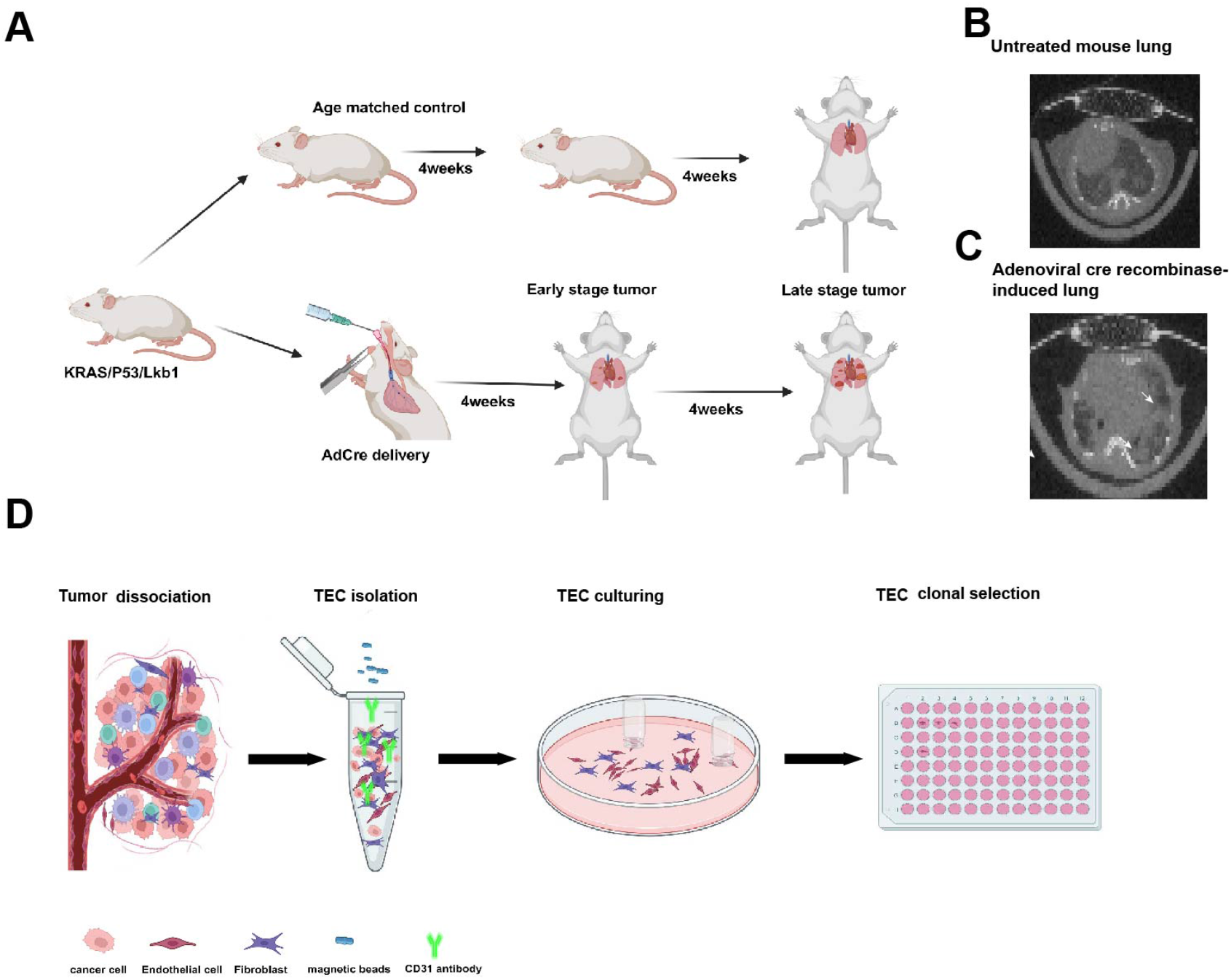
Lox-Stop-Lox (LSL)-KRAS^G12D^; Lkb1^L/L^; p53^L/L^ (KLP) mice were intranasally delivered 10^6^ PFUs of adenoviral Cre recombinase and monitored by *in vivo* CT imaging for tumor growth at both 4 weeks and 8 weeks (A), and lungs were visualized after 8 weeks by *in vivo* CT scanning for tumor excision. Panel B demonstrates no tumor growth in untreated mice, and panel C demonstrates lung tumors post-adenoviral Cre recombinase induction. (D) In the *in vitro* method, tumors are dissociated with collagenase and ECs were isolated using magnetic separation and a-CD31, cultured and clonally selected.

FACS was employed using a gating strategy for EPCAM, CD45, CD31 and LYVE1 to isolate ECs from lungs in both untreated mice and KLP mice (Figures 2A, i, ii and iii respectively). Separately, ECs were isolated using a dead/live sorting strategy in both untreated mice and tumor-bearing mice for LLC tumors and LN2A tumors (Figure 2B and 2C respectively) for live cells (Panels 2Bi and 2Ci) and the live cell population was further sorted for ZsG+ ECs (Panels 2Bii and 2Cii).

**Figure 2:**
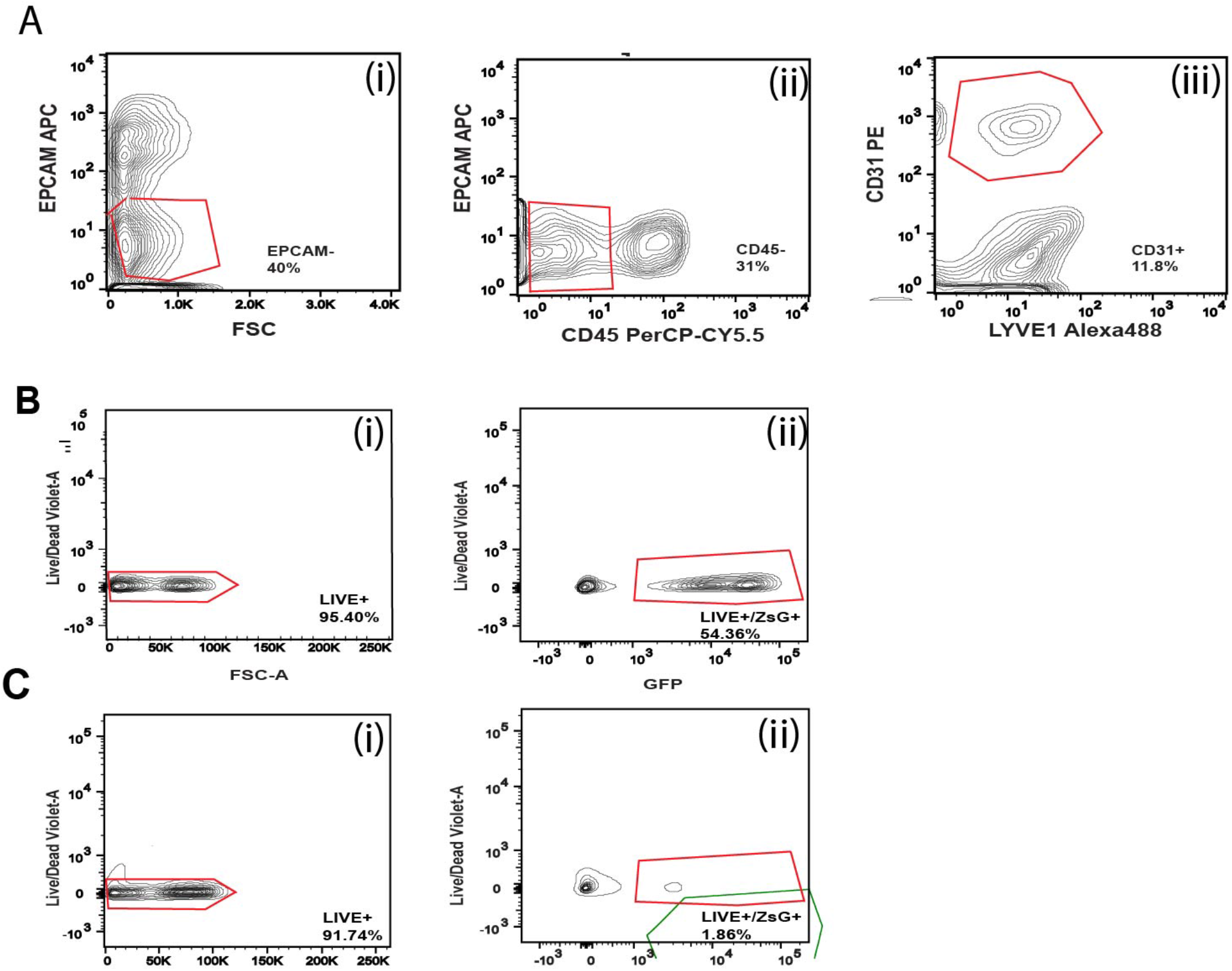
FACS was employed (A) using a gating strategy EPCAM (i), CD45 (ii), CD31 and LYVE1 (iii) to isolate ECs from lungs in both untreated mice and KLP mice. Separately, ECs were isolated using a dead/live sorting strategy in both untreated mice and tumor-bearing mice for LLC tumors (B) and LN2A tumors (C), with all live cells depicted (i) and ZsG+ ECs (ii).

### miRNA Expression Levels as Profiled by Nanostring

Using a Nanostring nCounter array system and ROSALIND, we evaluated in triplicate *in vivo* samples and observed that the global distribution of miRNAs is higher in NECs compared to TECs in the KLP mouse model, suggesting the overall expression profiles of miRNAs in TECs is significantly repressed, *P* <0.0001 (Figure 3A). In contrast, the *in vitro* cultured cell model traditionally used as a biomimetic system to study miRNA expression, the expression levels are uniform between NECs and TECs with a small but significant increase in miRNAs levels from TECs, *P* <0.0006 (Figure 3B). These data were followed by Volcano plot visualization to evaluate specific miRNAs in freshly isolated and cultured KLP ECs using NECs as a control. In the *in vivo* isolated TECs, miR-30b, miR-707 and miR-1981 were 2-3-fold significantly less expressed as compared with NECs (Figure 3C). Consistent with previously published reports on AngiomiRs, we also observed that miR-30b was significantly down-regulated in TECs^16^. Using the same approach with the *in vitro* EC profiles, miRNA-124, miR-200a and miR-186 were ~1.5-fold significantly less expressed in TECs than NECs, supported by previous literature demonstrating these miRNAs are down-regulated in pro-angiogenic states^7,17,18^. Conversely, miR-21 showed a 0.5-fold higher expression level, which is also in support of previous studies^19^. Taken together, these data suggest that the expression profiles of miRNAs substantially vary depending upon the methodology of isolation of TECs.

**Figure 3:**
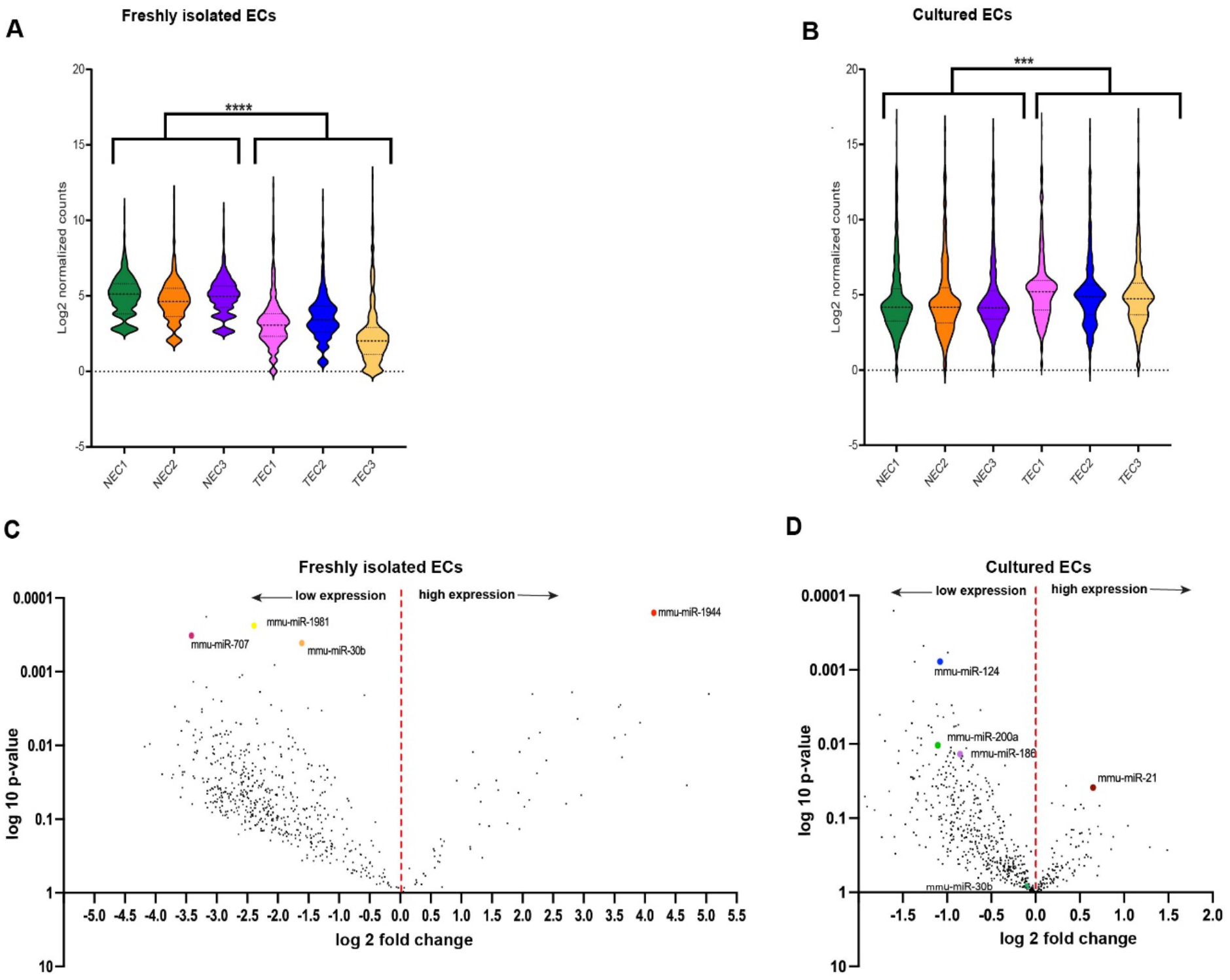
A violin plot was produced from KLP TECs and NECs that were freshly isolated (A) and cultured (B) to visualize the expression of miRNAs using a Nanostring nCounter array system and ROSALIND. Volcano plots evaluating specific miRNAs in freshly isolated (C) and cultured (D) KLP ECs. ^****^*P <* 0.0001, ^***^*P* < 0.001. Data are shown as the mean ± SEM incorporating biological and technical replicate samples. Two-tailed Student’s *t* test for 2-group comparisons.

### Validation of AngiomiR Expression Levels using qPCR

Next, to evaluate whether the *in vivo* and *in vitro* profiled AngiomiRs are generalizable to other cancer models, we utilized two highly metastatic lung cancer cell lines. Using tamoxifen-inducible (Cre-ERT2) VE-Cadherin (CDH5) LSL-ZsGreen reporter mice, following tamoxifen treatment, mice were orthotopically injected with firefly luciferase (fLuc) expressing cells (LLC-fLuc or LN2A-fLuc) (Figure 4A). This approach enabled non-invasive monitoring of disease progression, and once metastases developed tumors were harvested (Figure 4B and 4C). Following sorting, small RNAs were isolated from TECs and NECs freshly isolated from mice. Additionally, to evaluate the AngiomiR profiles in *in vitro* ECs, we obtained RNA from cultured TECs and NECs derived from KPL mice.

**Figure 4:**
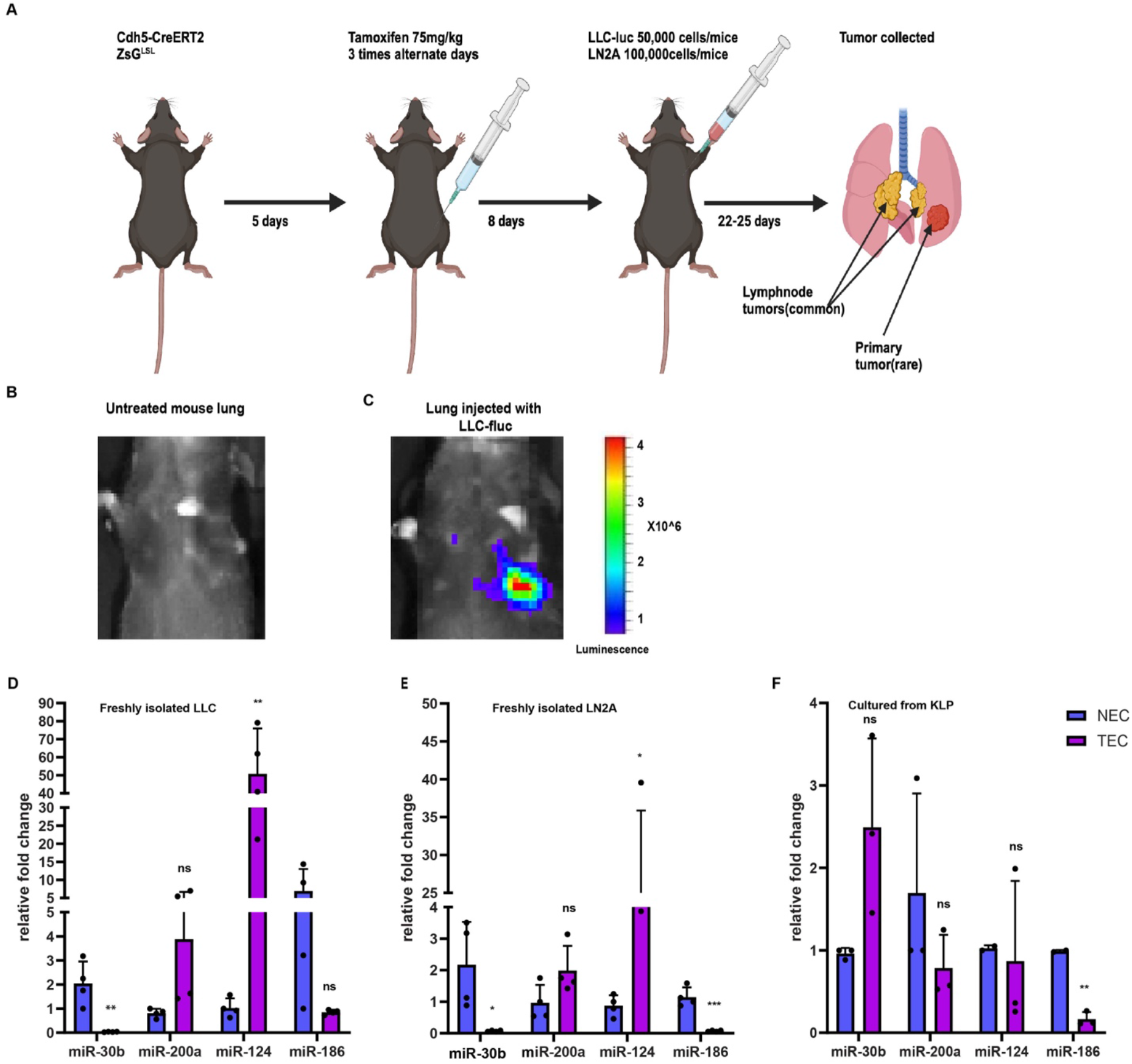
Tamoxifen-inducible (Cre-ERT2) VE-Cadherin (CDH5) LSL ZsGreen reporter mice were injected with tamoxifen and then injected with lung tumor cell lines conjugated with firefly luciferase (fLuc) (LLCfLuc and LN2AfLuc) (A) and resulting tumor volumes were measured by IVIS (B and C).Small RNAs were isolated from both NECs and TECs freshly isolated from mice injected with LLC-FLuc (D) and LN2A-Fluc (E) and from cultured KLP ECs (F) and then qPCR was performed to visualize miR-30B, miR-200A, miR124 and miRNA186. Relative fold changes were measured for miRNAs by normalizing values to NECs. ^***^*P <* 0.001, ^**^*P* < 0.01, ^*^*P* < 0.05. Data are shown as the mean ± SEM incorporating biological and technical replicate samples. Two-tailed Student’s *t* test for 2-group comparisons; 1-way ANOVA test for multiple comparisons.

Consistent with the AngiomiR profiles obtained from freshly isolated ECs (Figure 3C), in both LLC and LN2A models, we found TECs had significantly reduced miR-30b as compared with NECs (Figures 4D-E). We then evaluated ECs from these two models for the AngiomiR candidates obtained from the cultured TECs, and we only observed miR-186 as having significantly reduced expression in TECs as compared with NECs (Figures 4D-E). Conversely, when evaluating miRNA profiles in the cultured TECs and NECs, we found that miR-30b had a 2.5-fold higher expression, suggesting that cultured systems may not represent endogenous TEC miRNA expression. Similar to our findings in the LLC and LN2A freshly isolated TECs, we only observed miR-186 to have significantly reduced expression as compared with NECs (Figure 4F). Taken together, our findings demonstrate a substantial discordance in AngiomiR profiles based on the EC isolation method chosen. Furthermore, although we only evaluated a limited set of miRNAs, directly profiling freshly isolated ECs may yield more reproducible and rigorous results than that of cultured (*in vitro*) EC model systems.

## DISCUSSION

miRNAs are well understood regulators of cell homeostasis^20,21^. In epithelial cells, miRNAs have been implicated in the epithelial-to-mesenchymal transition (EMT), which increases tumor malignancy and invasiveness^22^. The miR-200 family is a classic suppressor of EMT and has also been found to block tumor angiogenesis through direct and indirect mechanisms by targeting interleukin-8 and CXCL1^7^. MiR-200a can facilitate the proliferation of cervical cancer cells and activates the HIF-1α/VEGF signaling pathway by targeting EGLN1, thus regulating VEGF mediated angiogenesis^23^. Interestingly, exosomes isolated from hepatocellular carcinoma (HCC) cell line culture over expressing miR-200b-3p inhibited endothelial ERG expression suggesting that decreased miR-200b-3p in cancer cells promotes angiogenesis in HCC tissues by enhancing endothelial ERG expression^24^. Also, several studies have found that miR-30b is downregulated in lung, breast, gastric and several other carcinomas through various regulatory mechanisms^25^. Earlier teams elucidated that miR-30b suppresses EMT in hepatoma cells and is significantly downregulated in hepatocellular carcinoma cells^26^. In non-small cell lung cancer studies, when downregulated, miR-30b expression in primary tumors inhibited proliferation and invasion by targeting EGFR^27,28^. Another study demonstrated that during capillary morphogenesis in HUVECS, miR-30b was downregulated under VEGF treatment. Overexpression of miR-30b by mimics activated capillary morphogenesis through enhanced expression of TGFβ2/Smad pathway^16^. These studies suggest that miRNAs can have both cancer cell intrinsic roles and effects on tumor angiogenesis.

Researchers have extensively used miRNA expression profiling from tissues or cultured cells. Computational methods of isolating miRNAs including learning tools and miRNA discovery tools, some of which use seed-matching, have been used to identify target sequences^29^. Researchers have used real-time qPCR methods with miR-specific RT primers as the gold standard method for miRNA profiling^30^. Research teams have produced hybridization-based high throughput microarray technology and combined this technology with methods like Northern blotting^31^. Microarrays can assess the expression of several miRNA at once using multiple probes. The assessment of miRNA levels can prove to be difficult, due to their dependence on the microenvironment, their short sequences, variations in unique expression, and the difficulty in designing specific probes^32^.

There is an inconsistency in the literature regarding miRNA expression levels depending on whether the cells were freshly isolated from tissue or by cell culture methods^10,11^. In a study examining the effect of acidic environments on miRNA levels, opposite changes in expression were seen for miR-7a in freshly isolated versus cultured cell lines^1^. A similar effect is seen in a study on mature glia, where some miRNAs that were not highly expressed in adult Mueller glia were shown to increase in cultured cells^11^. A similar contradictory result in the expression pattern of miRNAs appeared in hepatic stellate cells, where researchers used microarrays to examine the differential expression across isolated cells and cultured cells^3^. However, in some reports, consistent expression patterns may emerge through *in vivo* and *in vitro* methods. For example in colorectal cancer, a team observed that certain miRNA clusters that include miR-1, let-7, miR-15, miR-16, miR-99, miR-100, miR-125, miR-133, miR-143, miR-145, miR-192, miR-194, miR-195, miR-206, miR-215, miR-302, miR-367 and miR-497 are downregulated in both *in vivo* and *in vitro* isolation methods^5^. Whether the expression levels of angiogenic miRNAs show inconsistency based on the isolation methods thus became the major interest of this paper, after these previously published findings of discrepancies in the miRNA expression levels in *in vitro* and *in vivo* models.

In our study, we hypothesized that miRNA expression levels determined by cultured TECs (*in vitro* method) or freshly sorted TECs (*in vivo* method) would differ significantly from each other. The analysis of the Nanostring data using Rosalind revealed that the miRNA profiles are significantly different between freshly sorted NECs and TECs compared to cultured NECs and TECs. In our validation studies, discrepancies in miRNA expression were observed based on the lung cancer models used and isolation procedures. However, although we consistently observed significantly reduced miR-30b and miRNA-186 levels in all models, only miR-186 was observed to be reduced in cultured TECs. miR-30 family frequently have common targets as the members share an identical seed sequence. miR-30b and miR-30c targeted and down regulated DLL4, a ligand belonging to Notch signaling causing increased angiogenesis in vitro and in vivo^33^. In another study, miR-186-5p was found to be decreased in HUVECs in the context of hypoxia. Mechanistically miR-186-5p was found to be inhibited by TTTY15, a long non-coding RNA, and downregulation of miR-186-5p was rescued through TTTY15 silencing^34^.

In summary, our results demonstrate that miRNA expression levels in TECs vary depending on the isolation and culture methods. These findings suggest that even though feasible, miRNA profiling of cultured tumor endothelial cells may not fully recapitulate the *in vivo* profile of TECs. Considering the importance of miRNA expression and function in the regulation of the tumor microenvironment, we postulate that miRNA-based profiling of the tumor endothelium should be performed using freshly isolated ECs from tumors to most accurately capture their endogenous state.

## Acknowledgments

C.V.P. was supported in part by the National Institutes of Health (NIH) R01CA215075, R01CA258451, 1R01CA279532, 1R41CA246848, and 1R44CA284932, a Mentored Research Scholar Grants in Applied and Clinical Research (MRSG-14-222-01-RMC) from the American Cancer Society, the Jimmy V Foundation Scholar award, the University Cancer Research Fund, the Lung Cancer Research Foundation, and the Free to Breathe Metastasis Research Award. L.E. was in part supported by R01CA258451. S.H.A. was supported in part by a National Institute of General Medical Sciences under award 5T32 GM007092, NIH 1R41CA246848 and an NCBC TRG award. A.C.D. was in part supported by R01CA258451 and R01CA177875. The UNC Flow Cytometry Core Facility and Lineberger Comprehensive Cancer Center Animal Histopathology and Animal Studies Cores are all supported in part by a National Cancer Institute Center Core Support Grant (CA016086) to the UNC Lineberger Comprehensive Cancer Center. The UNC Flow Cytometry Core Facility is also supported in part by the North Carolina Biotech Center Institutional Support Grant 2012-IDG-1006.

## Author Contributions

Conception and design: C.V. Pecot, Development of methodology: L. Edatt, S.H. Azam, A.C. Dudley, C.V. Pecot, Acquisition of data (provided animals, performed experiments, provided facilities, etc.): L. Edatt, G. Dixit, S.H. Azam, A.C. Dudley, C.V. Pecot, Analysis and interpretation of data (e.g., statistical analysis, biostatistics, computational analysis): L. Edatt, G. Dixit, S.H. Azam, Y.S. Tsai, A.C. Dudley, C.V. Pecot, Writing, review, and/or revision of the manuscript: All authors, Administrative, technical, or material support (i.e., reporting or organizing data, constructing databases): L. Edatt, C.V. Pecot, Study supervision: C.V. Pecot

## REFERENCES

1 Kapora, E. et al. MicroRNA-505-5p functions as a tumor suppressor by targeting cyclin-dependent kinase 5 in cervical cancer. Biosci Rep 39 (2019). 10.1042/BSR20191221

2 Hippen, K. L., Loschi, M., Nicholls, J., MacDonald, K. P. A. & Blazar, B. R. Effects of MicroRNA on Regulatory T Cells and Implications for Adoptive Cellular Therapy to Ameliorate Graft-versus-Host Disease. Front Immunol 9, 57 (2018). 10.3389/fimmu.2018.00057

3 Li, Q. et al. MicroRNA-148a promotes apoptosis and suppresses growth of breast cancer cells by targeting B-cell lymphoma 2. Anticancer Drugs 28, 588–595 (2017). 10.1097/CAD.0000000000000498

4 Iqbal, M. A., Arora, S., Prakasam, G., Calin, G. A. & Syed, M. A. MicroRNA in lung cancer: role, mechanisms, pathways and therapeutic relevance. Mol Aspects Med 70, 3–20 (2019). 10.1016/j.mam.2018.07.003

5 Wang, D. et al. Exosome-encapsulated miRNAs contribute to CXCL12/CXCR4-induced liver metastasis of colorectal cancer by enhancing M2 polarization of macrophages. Cancer Lett 474, 36–52 (2020). 10.1016/j.canlet.2020.01.005

6 Orso, F. et al. Role of miRNAs in tumor and endothelial cell interactions during tumor progression. Semin Cancer Biol 60, 214–224 (2020). 10.1016/j.semcancer.2019.07.024

7 Pecot, C. V. et al. Tumour angiogenesis regulation by the miR-200 family. Nat Commun 4, 2427 (2013). 10.1038/ncomms3427

8 Korde, A. et al. Lung Endothelial MicroRNA-1 Regulates Tumor Growth and Angiogenesis. Am J Respir Crit Care Med 196, 1443–1455 (2017). 10.1164/rccm.201610-2157OC

9 Nikitenko, L. & Boshoff, C. Endothelial cells and cancer. Handb Exp Pharmacol, 307-334 (2006). 10.1007/3-540-36028-x_10

10 Soheilyfar, S. et al. In vivo and in vitro impact of miR-31 and miR-143 on the suppression of metastasis and invasion in breast cancer. J BUON 23, 1290–1296 (2018).

11 Wohl, S. G. & Reh, T. A. The microRNA expression profile of mouse Muller glia in vivo and in vitro. Sci Rep 6, 35423 (2016). 10.1038/srep35423

12 Kottakis, F. et al. LKB1 loss links serine metabolism to DNA methylation and tumorigenesis. Nature 539, 390–395 (2016). 10.1038/nature20132

13 Azam, S. H. et al. Quaking orchestrates a post-transcriptional regulatory network of endothelial cell cycle progression critical to angiogenesis and metastasis. Oncogene 38, 5191–5210 (2019). 10.1038/s41388-019-0786-6

14 McCann, J. V. et al. Endothelial miR-30c suppresses tumor growth via inhibition of TGF-beta-induced Serpine1. J Clin Invest 129, 1654–1670 (2019). 10.1172/JCI123106

15 Xiao, L., McCann, J. V. & Dudley, A. C. Isolation and Culture Expansion of Tumor-specific Endothelial Cells. J Vis Exp, e53072 (2015). 10.3791/53072

16 Howe, G. A., Kazda, K. & Addison, C. L. MicroRNA-30b controls endothelial cell capillary morphogenesis through regulation of transforming growth factor beta 2. PLoS One 12, e0185619 (2017). 10.1371/journal.pone.0185619

17 Shi, Z. et al. MiR-124 governs glioma growth and angiogenesis and enhances chemosensitivity by targeting R-Ras and N-Ras. Neuro Oncol 16, 1341–1353 (2014). 10.1093/neuonc/nou084

18 Becker, V. et al. Hypoxia-induced downregulation of microRNA-186-5p in endothelial cells promotes non-small cell lung cancer angiogenesis by upregulating protein kinase C alpha. Mol Ther Nucleic Acids 31, 421–436 (2023). 10.1016/j.omtn.2023.01.015

19 Sun, X. et al. Glioma stem cells-derived exosomes promote the angiogenic ability of endothelial cells through miR-21/VEGF signal. Oncotarget 8, 36137–36148 (2017). 10.18632/oncotarget.16661

20 Ofori, J. K. et al. Elevated miR-130a/miR130b/miR-152 expression reduces intracellular ATP levels in the pancreatic beta cell. Sci Rep 7, 44986 (2017). 10.1038/srep44986

21 Melkman-Zehavi, T. et al. miRNAs control insulin content in pancreatic beta-cells via downregulation of transcriptional repressors. EMBO J 30, 835–845 (2011). 10.1038/emboj.2010.361

22 Zhang, Q. et al. Roles and regulatory mechanisms of miR-30b in cancer, cardiovascular disease, and metabolic disorders (Review). Exp Ther Med 21, 44 (2021). 10.3892/etm.2020.9475

23 Su, X., Lang, C., Luan, A. & Zhao, P. MiR-200a promotes proliferation of cervical cancer cells by regulating HIF-1alpha/VEGF signaling pathway. J BUON 25, 1935–1940 (2020).

24 Wang, Y. et al. Exosomal delivery of miR-200b-3p suppresses the growth of hepatocellular carcinoma cells by targeting ERG- and VEGF-mediated angiogenesis. Gene 931, 148874 (2024). 10.1016/j.gene.2024.148874

25 Zhong, K., Chen, K., Han, L. & Li, B. MicroRNA-30b/c inhibits non-small cell lung cancer cell proliferation by targeting Rab18. BMC Cancer 14, 703 (2014). 10.1186/1471-2407-14-703

26 Dedeoglu, B. G. High-throughput approaches for microRNA expression analysis. Methods Mol Biol 1107, 91–103 (2014). 10.1007/978-1-62703-748-8_6

27 Riemann, A., Reime, S. & Thews, O. Acidic extracellular environment affects miRNA expression in tumors in vitro and in vivo. Int J Cancer 144, 1609–1618 (2019). 10.1002/ijc.31790

28 Maubach, G., Lim, M. C., Chen, J., Yang, H. & Zhuo, L. miRNA studies in in vitro and in vivo activated hepatic stellate cells. World J Gastroenterol 17, 2748–2773 (2011). 10.3748/wjg.v17.i22.

29 Pidikova, P., Reis, R. & Herichova, I. miRNA Clusters with Down-Regulated Expression in Human Colorectal Cancer and Their Regulation. Int J Mol Sci 21 (2020). 10.3390/ijms21134633

30 Li, L. et al. miR-139 and miR-200c regulate pancreatic cancer endothelial cell migration and angiogenesis. Oncol Rep 34, 51–58 (2015). 10.3892/or.2015.3945

31 Soufi-Zomorrod, M. et al. MicroRNAs modulating angiogenesis: miR-129-1 and miR-133 act as angio-miR in HUVECs. Tumour Biol 37, 9527–9534 (2016). 10.1007/s13277-016-4845-0

32 Zomorrod, M. S., Kouhkan, F., Soleimani, M., Aliyan, A. & Tasharrofi, N. Overexpression of miR-133 decrease primary endothelial cells proliferation and migration via FGFR1 targeting. Exp Cell Res 369, 11–16 (2018). 10.1016/j.yexcr.2018.02.020

33 Bridge, G. et al. The microRNA-30 family targets DLL4 to modulate endothelial cell behavior during angiogenesis. Blood 120, 5063–5072 (2012). 10.1182/blood-2012-04-423004

34 Zheng, J. et al. LncRNA TTTY15 regulates hypoxia-induced vascular endothelial cell injury via targeting miR-186-5p in cardiovascular disease. Eur Rev Med Pharmacol Sci 24, 3293–3301 (2020). 10.26355/eurrev_202003_20697

